# The non-thermogenic function of opossum UCP1 is independent of its cytoplasmic binding network

**DOI:** 10.1101/2025.04.27.650829

**Authors:** Giorgia Roticiani, Jürgen Kreiter, Elena E. Pohl

## Abstract

The primary function of uncoupling protein 1 (UCP1) is to mediate non-shivering thermogenesis in brown adipose tissue by facilitating proton transport across mitochondrial membranes. However, recent study has shown that UCP1 from the opossum (a marsupial) lacks thermogenic activity (Keipert et al., Science, 2024). This deficiency was attributed to two altered residues, K100 and Y289, within the cytosolic salt bridge network. Interestingly, the same residues are also present in UCP2 and UCP3 from murine species, both of which are known to transport protons in the presence of long-chain fatty acids (FAs). This raises questions about the validity of the proposed mechanism involving these residues in the loss of thermogenic function in opossum UCP1.

In this study, we performed conductance measurements of planar lipid bilayers reconstituted with either recombinant mouse UCP1, or UCP1 mutant (Q100K/F289Y), or UCP2, or UCP3. Our data demonstrate that the conductance of the UCP1 double mutant, UCP2, and UCP3 is comparable to that of wild-type mouse UCP1 in the presence of palmitic or arachidonic acids. These findings suggest that the altered cytosolic residues (K100 and Y289) do not explain the lack of thermogenic function in opossum UCP1. Thus, the underlying molecular mechanism responsible for the absence of thermogenesis in opossum UCP1 remains unresolved, and further studies are warranted to elucidate the precise cause of this dysfunction. Given the emerging interest in UCP1 uncoupling as a potential therapeutic approach for treating obesity by increasing energy expenditure, understanding the molecular basis of UCP1’s thermogenic dysfunction is of significant relevance.

## 1. INTRODUCTION

Mitochondrial uncoupling refers to the dissipation of the proton (H^+^) gradient across the inner mitochondrial membrane without the production of adenosine triphosphate (ATP). This process plays a crucial physiological role in non-shivering thermogenesis in brown adipose tissue (BAT) and is a physiologically relevant function for heat production in new-born animals, in cold-exposed rats and in arousing hibernators ^1^ Non-shivering thermogenesis is mediated by uncoupling protein 1 (UCP1), which is massively upregulated in response to cold exposure ^2^. For thermogenesis to occur, the presence of free fatty acids (FA) is essential; the protonophoric activity of UCP1 is inhibited by purine nucleotides (PN) ^3,4^.

Recently, it was reported that a UCP1 variant in the opossum (*Monodelphis domestica,* marsupial) lacks the ability to mediate non-shivering thermogenesis ^5^. Based on the comparative adipose tissue transcriptome and respirometry of HEK293 cells expressing UCP1 variants, it was proposed that amino acid residues glutamine Q100 and phenylalanine F289 play an essential role in the activity of UCP1, as they are replaced by lysine (K100) and tyrosine (Y289) in the sequence of opossum UCP1 (Fig. 1A, Fig. S1). These residues are located at the cytosolic ends of helix 2 and 6 and are involved in the cytosolic salt bridge network (Fig. 1B). However, K100 and Y289 are also present in the UCP1 homologs UCP2 and UCP3, which are also known to mediate H^+^ transport in the presence of FA in mitochondria. UCP2 and UCP3 were shown to transport H^+^ with transport rates similar for UCP1, (reviewed in ^6^).

**Figure 1.**
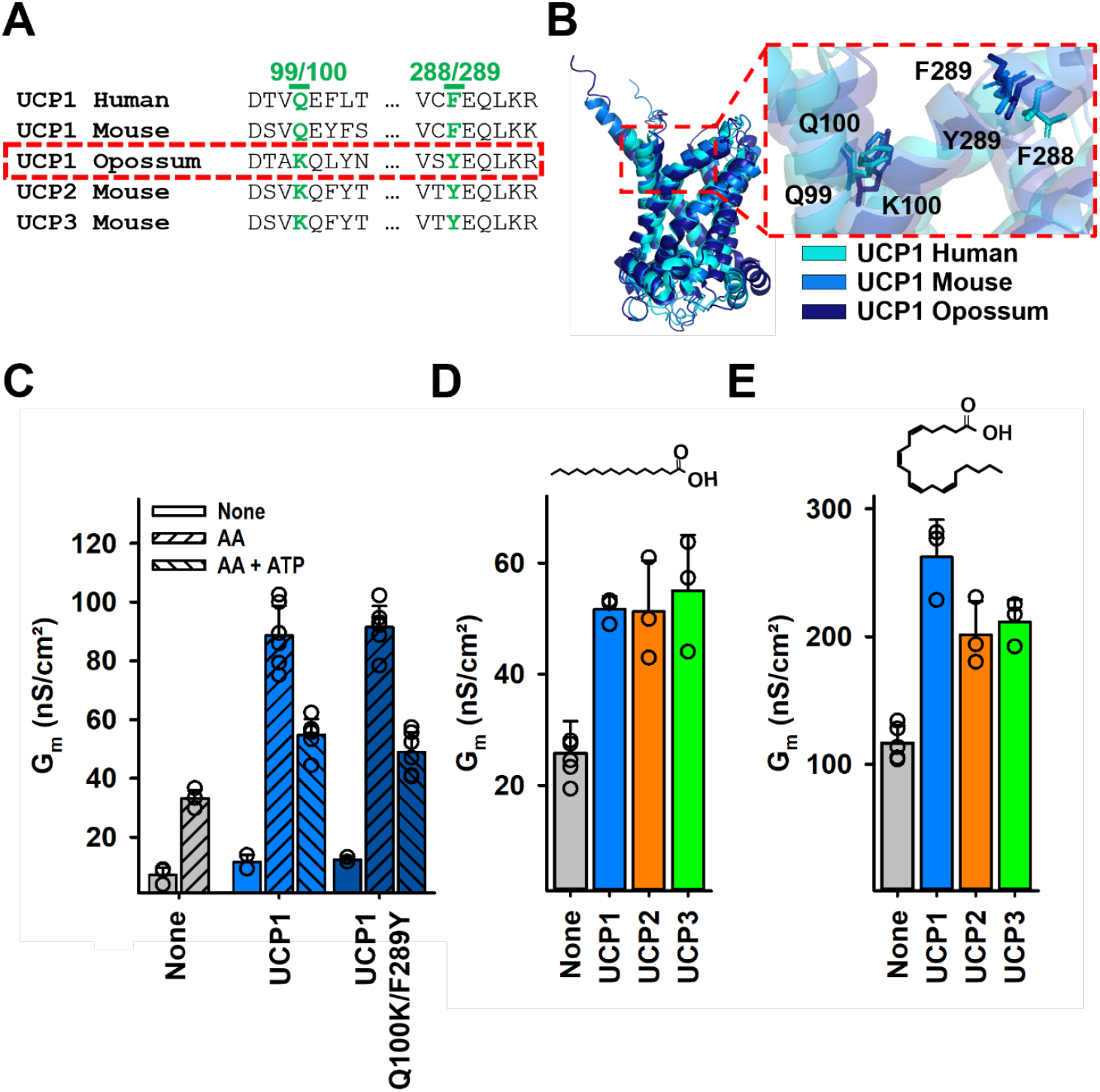
Proton transport mediated by UCP1, UCP1-Q100K/F289Y, UCP2 and UCP3 in the presence of palmitic or arachidonic acid. (A) Sequence alignment of human, mouse and opossum UCP1 and mouse UCP2 and UCP3 around residues 100 and 289. The sequence of opossum UCP1 is highlighted by the red box and the residues of interest in green, respectively. (B) Superposition of the structures of human, mouse and opossum UCP1 generated in AlphaFold based on the structure of human UCP1 (PDB: 8HBV). The box indicates the orientation of the residues of interest. (C) Total membrane conductance (G_m_) of lipid bilayers reconstituted with mUCP1, mUCP1 - Q100K/F289Y, and no protein in the absence of AA (first bar in each set), in the presence of AA (second bar in each set), and in the presence of AA and 4 mM ATP (third bar in sets 2 and 3). (D-E) Total membrane conductance (G_m_) of lipid bilayers reconstituted with palmitic acid (D) or arachidonic acid (E) in the absence of UCPs (gray) and in the presence of UCP1 (blue), UCP2 (orange) or UCP3 (green) at 170 mV. Bars represent the mean and standard deviation of at least 3 independent measurements. In (C-E), the data points represent the signal value of each experiment. For all measurements, lipid bilayers were prepared from 45:45:10 mol% DOPC:DOPE:CL, reconstituted with 15 mol% fatty acid where indicated. The lipid concentration was 1.5 mg/mL and the protein concentration was 4 μg/(mg lipid). Measurements were performed in assay buffer containing 50 mM Na_2_SO_4_, 10 mM Tris, 10 mM MES, and 0.6 mM EGTA at pH = 7.34 and T = 306 K.

Additionally, the putative involvement of the cytosolic salt bridge in proton transport contradicts the FA-cycling model in which H^+^ transport is achieved by protonation/deprotonation of FA at the membrane surface ^7^. This predicts the binding of FA to the matrix side of UCP1. Mutation of residue R60 of UCP2 and R59 of AAC1 on the matrix side significantly decreased H^+^ transport ^8, 9^. However, the positively charged K67 on the matrix side is present in both mUCP1 and opossum UCP1, which contrasts with their apparently different activity.

Taken together, the above-mentioned observations cast doubt on the idea that the amino acid replacement of Q100 and F289 by K100 and Y289 in opossum UCP1 accounts for the loss of its protonophoric function.

In this study, we used site-directed mutagenesis and planar bilayer conductance recordings to directly compare H^+^ transport mediated by recombinant mouse UCP1 and the UCP1 double mutant (Q100K/F289Y) in the presence of arachidonic acid (AA). Furthermore, we analyzed the activation of mUCP1, mUCP2 and mUCP3 by palmitic acid (PA) and AA at potentials up to 170 mV, corresponding to different functional states of mitochondria.

## 2. RESULTS AND DISCUSSION

We first constructed the mUCP1 mutant, replacing Q100 and F289 with K100 and Y289, and compared the conductance, *G*_*m*_, of membranes reconstituted with wt and mutant UCP1 (Fig. 1C, Fig. S2). As expected, both variants showed no H^+^ transport in the absence of arachidonic acid (AA). Addition of AA increased G_m_ in both mUCP1 and opossum-like mUCP1. Also, the inhibition (approx. 60-70 %) of the wt and mutant mUCP1 by ATP was similar, demonstrating that the substrate binding site is maintained and ensures the structural integrity of mutant UCP1.

The assumption that Q100 and F289 of eutherian UCP1 are essential for FA-activated H^+^ transport implies, in principle, that activation by FA is a result of FA-UCP1 interaction from the cytosolic side. However, since the mutation of Q100 and F289 did not affect the activation of UCP1 by FA, it can at least be assumed that these residues are not involved in FA binding from the cytosolic side.

We have previously proposed that the FA anion (FA^−^) binds to the mUCP1 and ADP/ATP carrier from the matrix side and slides along the protein-lipid interface ^10^. It is similar to the mechanism of lipid flip-flop mediated by lipid scramblases ^11^. Since the mechanism involves residues that are highly conserved within the SLC25 family ^12,13^, the mechanism of FA^−^ transport may also be valid for UCP2 and UCP3.

We therefore measured the conductance of membranes reconstituted with mUCP1 homologs, mUCP2 and mUCP3, since their sequence contains the same residues (K100 and Y289) as opossum UCP1. We obtained current-voltage recordings at applied voltages up to 220 mV (Fig. S3) in the presence of saturated palmitic acid (PA, C16:0) and polyunsaturated arachidonic acid (AA, C20:4). For both FAs, G_m_ was significantly higher in the presence of UCP1-3 than in lipid bilayers reconstituted with PA (Fig. 1D) or AA (Fig. 1E) alone. As expected, G_m_ was 3-to 5-fold higher in the presence of AA than PA.

In this work, we have shown that residues K100 and Y289 in opossum UCP1 do not alter the activity of UCP1. The mUCP1-Q100K/F289Y mutant showed no difference in H^+^ transport compared to mUCP1 in the presence of AA. Thus, our results challenge the assumptions that were drawn by the phylogeny and respirometry of UCP1 variants by Keipert et al. ^5^. Further evidence comes from the activity of mUCP2 and mUCP3, which contain the same residues K100 and Y289 as opossum UCP1 and have also been shown to mediate H^+^ transport in the presence of FA. The exact reason why opossum UCP1 fails to mediate non-shivering thermogenesis remains unclear and requires more careful analysis of additional amino acids that are different in its sequence.

## MATERIALS AND METHODS

### Chemicals

Sodium sulfate (Na_2_SO_4_), 2-(N-morpholino) ethanesulfonic acid (MES), tris-(hydroxymethyl)-aminomethane (Tris), sodium dodecyl sulfate (SDS) and ethylene glycol-bis (β-aminoethyl ether)-N,N,N’,N’-tetraacetic acid (EGTA) were purchased from Carl Roth GmbH & Co. K.G. (Karlsruhe, Germany). Hexane, hexadecane, adenosine 5’-triphosphate (ATP), guanosine 5’ -triphosphate (GTP) and agarose were purchased from Sigma-Aldrich Co. (Vienna, Austria). 1,2-dioleoyl-sn-glycero-3-phosphocholine (DOPC), 1,2-dioleoyl-sn-glycero-3-phosphoethanolamine (DOPE) and cardiolipin (CL) were obtained from Avanti Polar Lipids Inc (Alabaster AL, USA) and chloroform from Sanova Pharma GesmbH (Vienna, Austria). Arachidonic acid (AA, C20:4) was purchased from Larodan (Biozol, Eching, Germany) and palmitic acid (PA, C16:0) from Sigma-Aldrich Co. (Vienna, Austria).

### Production of UCP1 mutant, isolation, purification and reconstitution of murine UCP1, UCP2 and UCP3 into liposomes

To generate the UCP1 mutant, we carried out site-directed mutagenesis of Q100 to K and F289 to Y using the Q5 Site-Directed Mutagenesis Kit (New England Biolabs, Austria). The sequence was verified by both DNA and amino acid sequencing. The refolding, purification and reconstitution of wild-type murine UCPs (mUCP1, mUCP2 and mUCP3) and the mUCP1-Q100K/F289Y double mutant was performed as previously described ^14,15^. Both proteins were reconstituted in parallel to minimize reconstitution artifacts of mutant mUCP1 and protein purity was verified by SDS-PAGE and silver staining. Protein concentration was determined using the Micro BCA Protein Assay Kit (Thermo Fisher Scientific, Waltham, MA, USA). The following protein batches were used in this study: mUCP1 – 148, 150; mUCP1-Q100K/F289Y – 2, 3; mUCP2 – 59, 61 and mUCP3 – 42, 43.

### Measurements of total membrane conductance in planar lipid bilayers

Planar lipid bilayers were formed on the tip of a disposable plastic pipette ^3^. Membrane formation was verified by membrane capacitance measurements (C = 0.75 ± 0.03 μF/cm^2^) and is independent of protein content, fatty acid (FA) and inhibitors. Current-voltage (I-U) recordings were performed at T = 306 K and recorded with a patch-clamp amplifier (EPC 10, HEKA Elektronik, Dr. Schulze GmbH, Lambrecht, Germany) at an acquisition rate of 510 Hz. Total membrane conductance (G_m_) at 0 mV was obtained from the slope of a linear fit to the current from a voltage ramp from − 50 mV to + 50 mV. G_m_ at 170 mV was calculated from an exponential fit to I-U recordings from a voltage ramp from 0 – 220 mV ^14^ and is identical to a linear fit performed to data in the range of 160 to 180 mV. Proteoliposomes were solved in assay buffer solution (50 mM Na_2_SO_4_, 10 mM Tris, 10 mM MES and 0.6 mM EGTA, pH = 7.34). Fatty acids AA or PA added to the lipids while dissolved chloroform. ATP was dissolved in assay buffer solution and added to the proteoliposomes in buffer solution.

### Statistics

Data are presented as the mean @ standard deviation of at least three independent experiments and at least two independent refoldings and reconstitutions of the proteins. Individual data points are shown in the figures.

## ACKNOWLEDGMENTS

We thank Sarah Bardakji for the help in generating the mUCP1-Q100K/F289Q mutation and Alina Pashkovskaya for the help in the refolding, purification and reconstitution of the proteins into liposomes.

## AUTHOR CONTRIBUTIONS

E.P., G.R. and J.K. designed the experiments. G.R. and J.K. conducted the experiments, interpreted the data and prepared the figures. All authors discussed the results and edited and revised the manuscript.

## SUPPLEMENTARY MATERIALS

Material and Methods

Fig. S1 – Fig. S3

## DATA AND MATERIALS AVAILABILITY

All data are available in the main article or the supplementary materials and from the corresponding author upon reasonable request.

