## Supplementary Figures for "The non-thermogenic function of opossum UCP1 is independent of its cytoplasmic binding network"

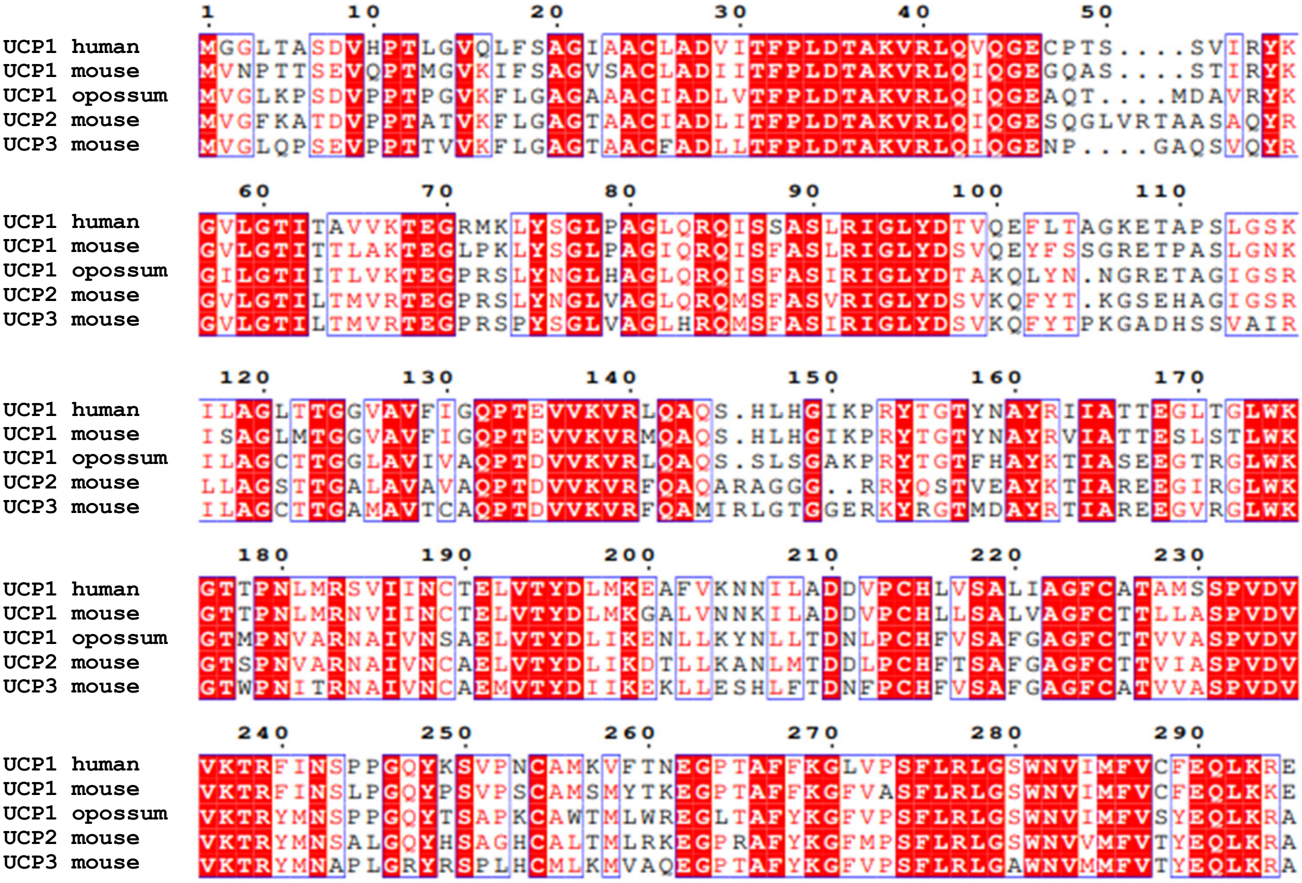


**Figure S1.** Alignment of the full amino acid sequences of human UCP1 (UniProt ID:P25874), mouse UCP1 (P12242), opossum UCP1 (F6X9Y8), mouse UCP2 (P70406) and mouse UCP3 (P56501).

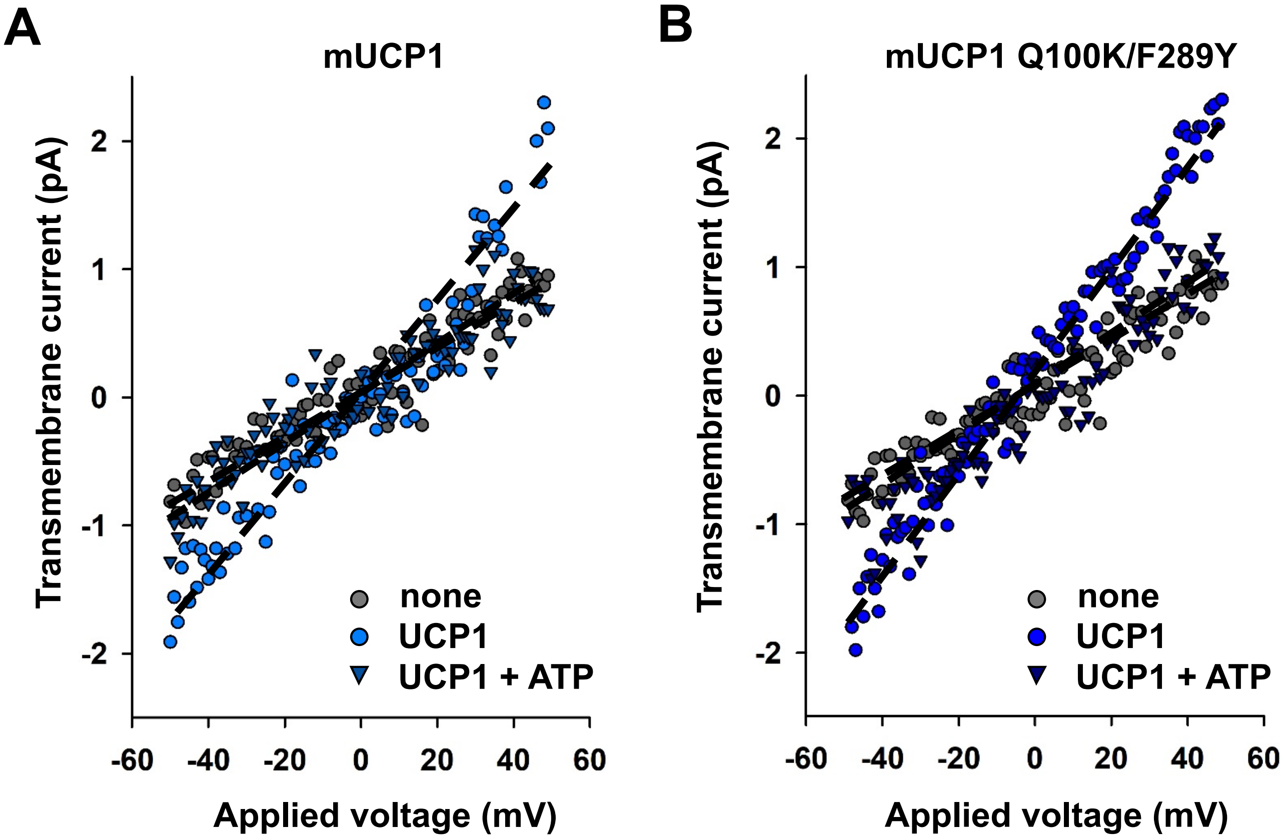


**Figure S2.** Current-voltage recordings of lipid bilayers reconstituted with 15 mol% AA in the absence (gray circles) and in the presence (blue circles) of mUCP1 (A) or mUCP1-Q100K/F289Y (B) and in the presence of UCP1 and 4 mM ATP (blue triangles). Black dashed lines represent are a linear fit to the data. Experimental conditions were as described in Figure 1.

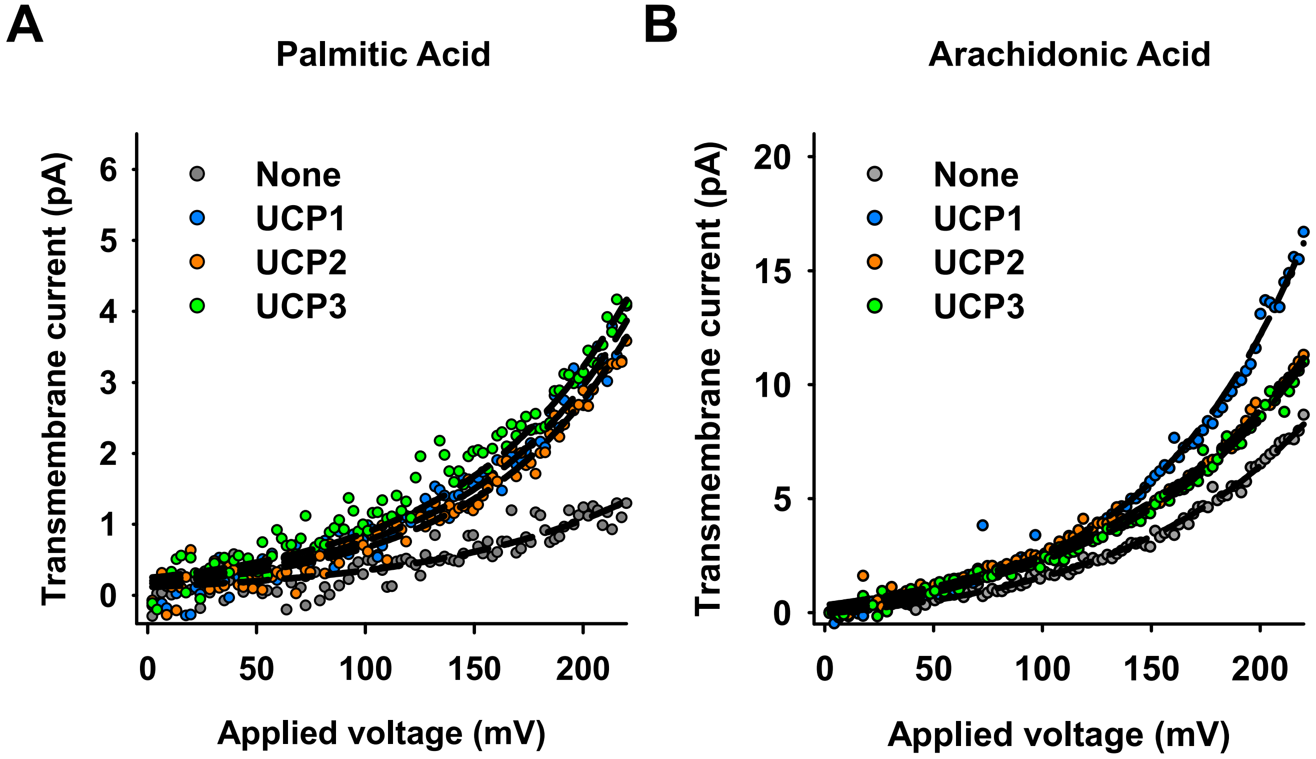


**Figure S3.** Current-voltage recordings of lipid bilayers reconstituted with 15 mol% palmitic acid (PA, A) or arachidonic acid (AA, B) in the absence of UCPs (gray) and in the presence of mUCP1 (blue), mUCP2 (orange) and mUCP3 (green). Black dashed lines represent a least-squares fit of an exponential function to the data. Experimental conditions were as described in Figure 1.
